# Induction of microbial oxidative stress as a new strategy to enhance the enzymatic degradation of organic micropollutants in wastewater

**DOI:** 10.1101/544205

**Authors:** Amrita Bains, Octavio Perez-Garcia, Gavin Lear, David Greenwood, Simon Swift, Martin Middleditch, Edward Kolodziej, Naresh Singhal

## Abstract

Organic micropollutants (OMPs) are pervasive anthropogenic contaminants of fresh and marine waters with known potential to adversely affect aquatic life (e.g. endocrine disruption). Their ubiquitous environmental occurrence is primarily attributed to wastewater treatment plant discharges following their incomplete removal by common biological treatment processes. This study assesses a new strategy for promoting the degradation of six model OMPs (i.e. sulfamethoxazole, carbamazepine, tylosin, atrazine, naproxen and ibuprofen) by stimulating microbial oxidoreductase production to counter the effects of oxidative stress caused by oxygen perturbation. Microbial cultures from dairy farm wastewater were exposed to a cyclical ON-OFF perturbations of oxygen supply, ranging from 0.16 to 2 cycles per hour (i.e. 2, 1, 0.5, 0.25 and 0.16 cycles/hour), in laboratory bioreactors. The activity and relative abundances of microbial oxidoreductases (such as peroxidases, cytochromes P450) were upregulated by oxygen perturbation. In comparison to controls subjected to constant oxygen levels, OMP concentrations in perturbed cultures decreased by 70±9% (mean ± SD). A distance-based linear model confirmed strong positive correlations between the relative abundance of the bacterial families, *Rhodocyclaceae, Syntrophaceae* and *Syntrophobacteraceae,* and oxygen perturbations. Our results confirm that intentional perturbation of oxygen supply to induce microbial oxidative stress can improve OMP removal efficiencies in wastewater treatment bioreactors.

## Significance

Oxygen concentration is a critical control variable for biological processes in conventional wastewater treatment systems. We provide a simple approach that could be used in existing wastewater treatment infrastructure to enhance removals of otherwise persistent OMPs by exploiting oxidative stress responses of microbes. Our study illustrates the ability of fluctuating oxygen conditions to affect microbial activity and polulation by upregulating oxidoreductases linked to the removal of organic micropollutants. The incorporation of inexpensive and easy to implement dissolved oxygen supply strategies for more efficient OMPs biotransformation provides a realistic approach to improving the treatment performance of biological processes.

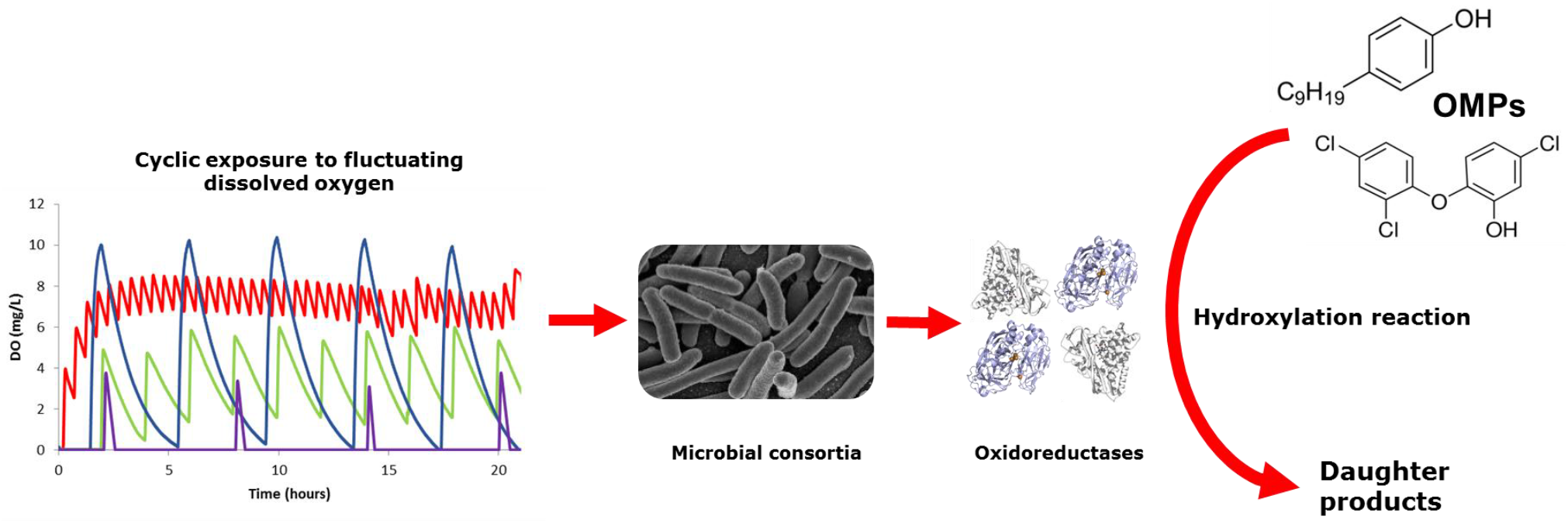

## Introduction

The discharge of trace levels of pharmaceuticals, industrial chemicals, pesticides and personal care products into aquatic environments is an issue of global environmental concern (1–3). Because of their resistance to conventional wastewater treatment and environmental persistence as complex mixtures, these organic micropollutants (OMPs) present risks of adverse ecological impact to biota via direct toxicity and endocrine disruption (4–6). For example, occurrence of the herbicide, atrazine, even at concentrations below 6 μg/L in receiving waters, caused behavioural changes in the zebrafish, *Brachydanio rerio,* such as a preference for dark habitats and alterations in swimming due to failure of its nervous system (7). OMP exposure effects may be additive; for example studies by (8) and (9) have reported that exposure of female *Danio rerio* to pharmaceutical mixtures of acetaminophen, carbamazepine, gemfibrozil and venlafaxine significantly reduces fecundity, and disproportionate to effects expected for individual exposures. Such concerns about the environmental occurrence of OMPs have triggered attempts to enhance their biodegradation during wastewater treatment (10). Although conventional activated sludge methodologies are not specifically optimised to eliminate OMPs (11), exploiting the diverse enzymatic potential of microbial consortia has been identified as a critical pathway for OMP removal (12–14).

In the period since the first oxidative conditions arose on Earth due to early photosynthesis, microbes have evolved to survive exposure to a variety of harmful oxidative stresses (15). For instance, oxidative stress generated by the high concentration of intracellular reactive oxygen species (ROS) including superoxide (O_2_̄), hydrogen peroxide (H_2_O_2_) and hydroxyl radicals (OH·), and *ipso facto*, induces oxidative enzyme gene expression (16) and consequent synthesis of antioxidative enzymes such as peroxidases and oxidoreductases to protect against such oxidative stress (17, 18). From a wastewater treatment perspective, intentional manipulations of environmental oxygen concentrations could stimulate the production of these enzymes, some of which are also capable of degrading toxic substances including polycyclic aromatic hydrocarbons and organophosphorus contaminants (19–23). Therefore, oxygen perturbations might pre-dispose cells, through transcriptional responses of their oxidoreductases to protect against iterative oxidative stress, to enhance OMP degradation. This proposed mechanism implies that simple oxygen control strategies within wastewater treatment bioreactors may provide a cost-effective solution for enhanced biological OMP removal.

In this study we investigated the impact of dynamic variation in oxygen concentration on OMP biotransformation by representative wastewater microbial consortia. We hypothesise that varying dissolved oxygen concentration can enhance the degradation of OMPs by inducing oxidative stress and forcing microbes to produce oxidoreductases. Specifically, we assessed the effect of varying frequency of oxygen perturbations on microbial consortia by monitoring the synthesis of oxidative biocatalysts (oxidoreductases) along with the removal of OMPs.

## Results

### Influence of oxygen perturbations on microbial enzymatic activity

Five frequencies (2, 1, 0.5, 0.25 and 0.16 cycles per hour) were tested at two aeration regimes (high-DO, up to 10 mg-DO/L and low-DO, up to 4 mg-DO/L) that generated cyclic patterns of dissolved oxygen concentrations in bioreactors (Figure 1, Table 1). The oxidoreductase (lignin peroxidase, horseradish peroxidase, laccase and cytochrome P450) and hydrolase (beta-glucosaminidase, beta-glucosidase) activities in bioreactors were detected spectrophotometrically by the oxidation of their respective chromogenic substrates (Methylene Blue [MB], Azure B [AB], 3,4-Dihydroxy-L-phenylalanine [L-DOPA], 2,2’-Azino-bis-(3-ethylbenzothiazoline-6-sulfonic acid)[ABTS], Sudan Orange G [SO], 4-nitrophenyl-dodecanoate [pNP-12], Indole and 4-Aminoantipyrine [4-AAP], 4-nitrophenyl N-acetyl-β-D-glucosaminide [pNP-A] and 4-nitrophenyl-β-D-glucopyranoside [pNP-G] (Table S1). Cultures perturbed in high-DO regimes at 0.25 cycles/hr showed significantly higher (ANOVA, p < 0.05) beta-glucosidase (pNP-G), lignin peroxidase (AB, MB) and horseradish peroxidase (ABTS) activity compared to cultures treated at 1 (lignin peroxidase (AB, MB), beta-glucosidase (pNP-G) and cytochrome P450 (Indole)) or 2 (lignin peroxidase (AB, MB) only) cycles/hr. The cultures exposed to 0.25 and 0.5 cycles/hr and maintained within a low-DO range showed high (ANOVA, p < 0.05) cytochrome P450 (Indole, 4-AAP), lignin peroxidase (AB, MB), and horseradish peroxidase (L-DOPA, ABTS) activity whereas 0.16 cycles/hr subjected cultures exhibited more beta-glucosaminidase (pNP-A), beta-glucosidase (pNP-G), lignin peroxidase (MB, AB) and horseradish peroxidase (L-DOPA, ABTS) activity (Figure 2a).

**Table 1.** Dissolved oxygen (DO) conditions used for the OMP removal studies. Controls had constant (unperturbed; 0 cycles/hr) DO supply at either high or low concentrations.

| Operational condition | Dissolved Oxygen range (mg/L) | Perturbation frequency (cycles/hr) | Dissolved Oxygen shock durations (min.) |  |
| --- | --- | --- | --- | --- |
|  |  |  | Oxygenation duration (min.) | Deoxygenation duration (min.) |
| High-DO Aerobic | 8-9 | 0 | Constant | Constant |
|  | 0-10 | 0.25 | 30 | 210 |
|  | 4-6 | 1 | 10 | 50 |
|  | 6-8 | 2 | 8 | 22 |
| Low-DO Aerobic | 2-3 | 0 | Constant | Constant |
|  | 0-3 | 0.16 | 6 | 354 |
|  | 0-4 | 0.25 | 6 | 234 |
|  | 2-5 | 0.5 | 12 | 106 |

**Figure 1.**
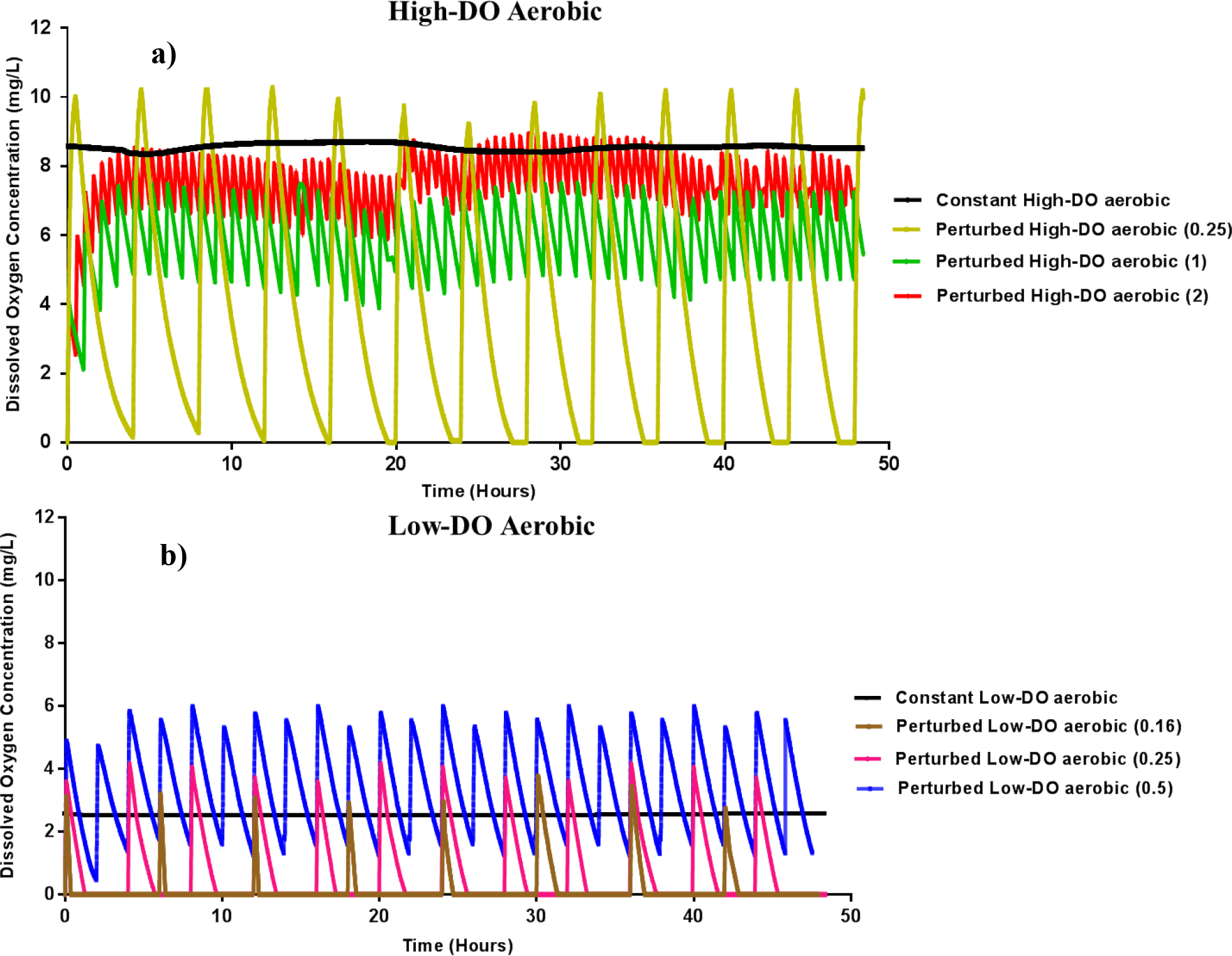
Dissolved oxygen profiles of microbial mixed cultures under: (a) high-DO and (b) low DO perturbation in comparison to non-perturbed conditions (constant). Stable DO profiles were achieved in the reactors that were provided constant aerobic (high and low) DO conditions, while aerobic-anaerobic DO transitions were induced with perturbed conditions.

**Figure 2.**
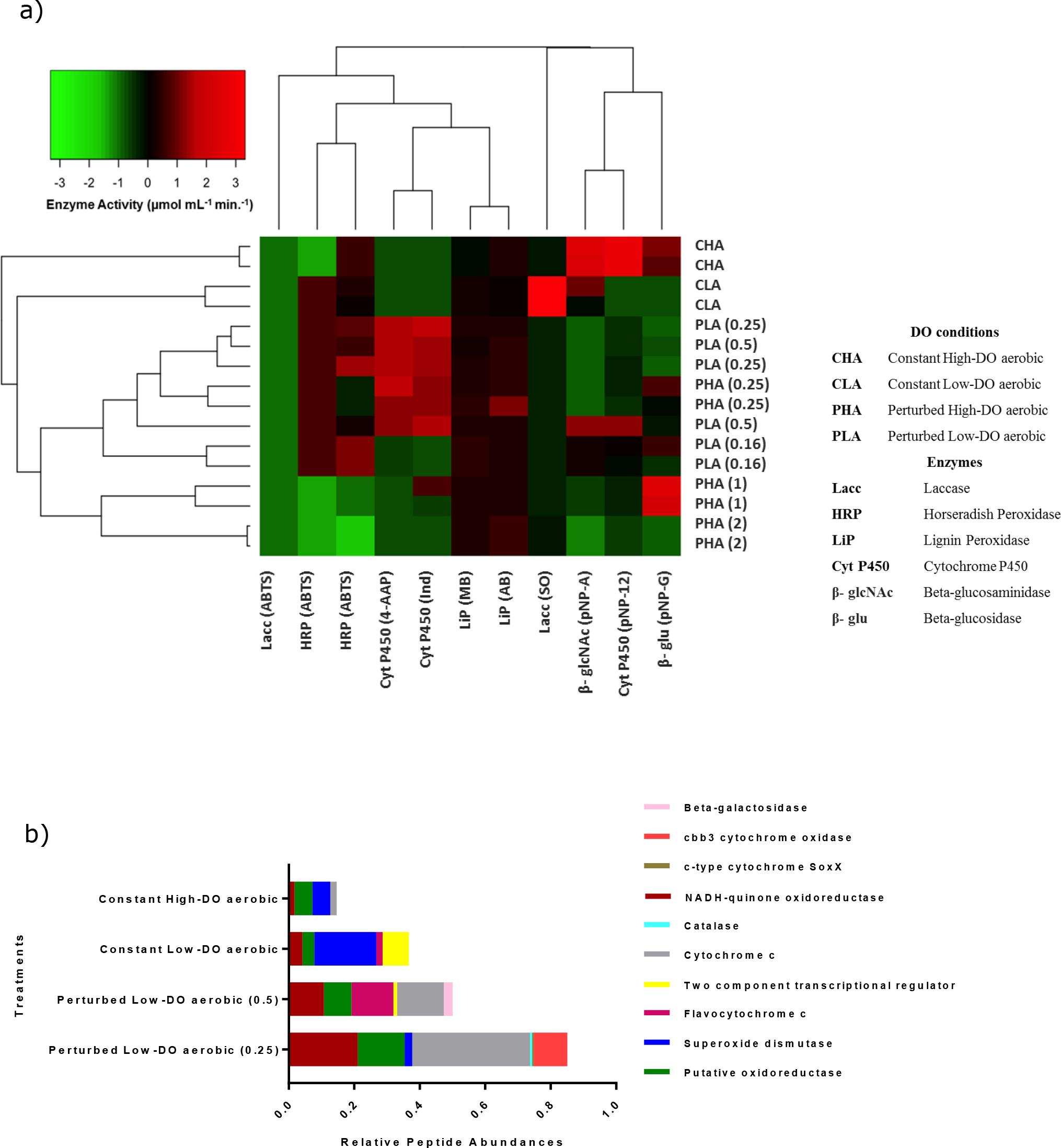
(a) Oxidoreductase activities in: (a) DO perturbed and non-perturbed samples, and (b) relative peptide abundances of expressed oxidoreductases (95 % confidence interval). Different rows represent DO perturbed frequencies in high and low-DO regime; different columns represent expressed oxidoreductase activities. The color intensity in each panel shows the auto-scaled enzyme activity (μmol mL^−1^ min.^−1^) (a) across the gradient from red (highest enzyme activity) to green (lowest enzyme activity).

By contrast, under constant non-perturbed oxygen concentrations, the high-DO aerobic cultures exhibited greater (ANOVA, p < 0.05) cytochrome P450 (pNP-12), beta-glucosaminidase (pNP-A), beta-glucosidase (pNP-G), lignin peroxidase (AB) and horseradish peroxidase (L-DOPA) activity, while the low-DO aerobic samples showed greater laccase (SO), beta-glucosaminidase (pNP-A), lignin peroxidase (MB) and horseradish peroxidase (ABTS, L-DOPA) activity. Proteomic analysis supported the above enzyme activity profiles with significant (95% confidence interval) observations showing that maximum relative abundance of expressed oxidoreductases (cytochromes, peroxidases and oxidases) had occurred in perturbed low-DO cultures (0.5 and 0.25 cycles/hr) (Figure 2b).

### Micropollutant removal by mixed cultures

The removal of OMPs (at 0.1 mg/L initial nominal concentrations) was significantly (ANOVA, p < 0.05) increased by the perturbed DO conditions (removing OMPs below the limit of quantification (LOQ)). The perturbed treatments showed 70±9% (mean ± SD) removal of sulfamethoxazole, carbamazepine, tylosin, atrazine, naproxen and ibuprofen, with an average residual concentration of 0.04 mg/L for individual OMPs remaining at the end of the experiments. Cultures exposed to 0.5 cycles/hr in the low-DO regime exhibited near log-removal (~0.01-0.02 mg/L residual concentrations) of sulfamethoxazole, naproxen and ibuprofen, whereas carbamazepine, tylosin and atrazine concentrations in the cultures subjected to 0.16 and 0.25 cycles/hr in low-DO range were attenuated to residuals of 0.012-0.04 mg/L (Figure 3). Cultures under constant low-DO conditions showed 35±8% removal of all compounds compared to constant high-DO treated cultures with 25±6% removal. On average, 0.06 mg/L residual concentrations of OMPs were detected in constant low-DO cultures while constant high-DO cultures exhibited 0.08 mg/L residual concentration (Figure 3). There were no significant differences (post-hoc Tukey test, p > 0.05) for sulfamethoxazole, carbamazepine and tylosin between DO perturbed and non-perturbed constant high and low-DO aerobic cultures. However, a significant removal (post-hoc Tukey test, p < 0.05) of atrazine, naproxen and ibuprofen can be seen under low-DO aerobic perturbed cultures compared to high-DO aerobic and non-perturbed constant cultures. Autoclaved biomass, used as a negative control, showed a loss of 15±6% for all OMPs (data not shown) which is attributed to abiotic factors such as hydrophobic partitioning and sorption to the sludge solids. Figure S1 and Tables S2 and S3 in the supplementary information show OMP removal, OMP residual concentrations and quality control measures, respectively.

**Figure 3.**
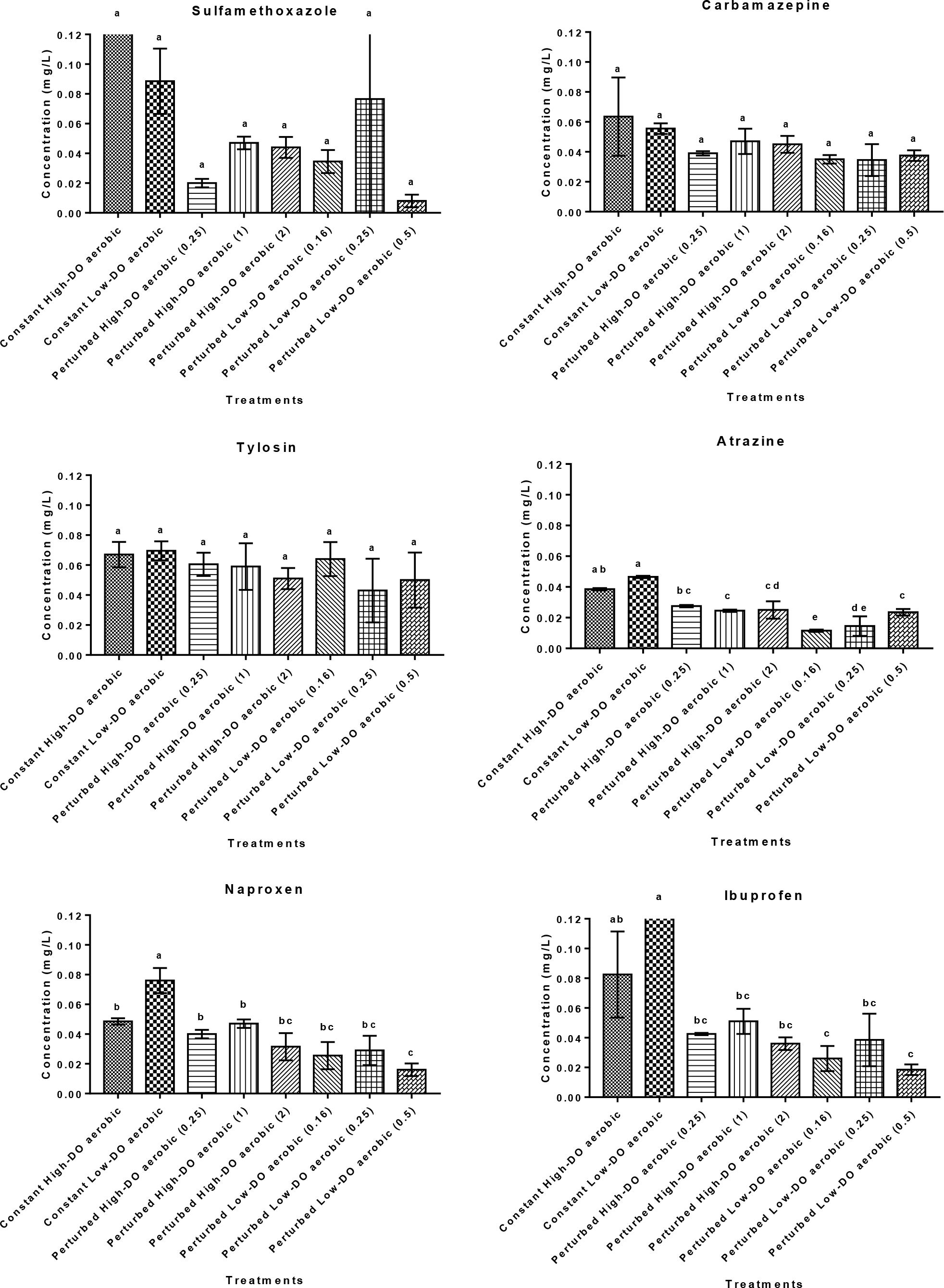
Residual concentrations of OMPs in supply-perturbed or constant supply (control) aerobic cultures at either high (8-9 mg/L) or low (2-3 mg/L) DO concentrations. Different letters above bars in the same graph indicate significant statistical differences between datasets via ANOVA (p = 0.05) and post hoc Tukey tests; bars indicated with the same letter are not significantly different

### Influence of oxidative stress on microbial speciation

A total of 527 operational taxonomic units (97% OTUs) were identified in the cultures, of which 513 represented Bacteria and 13 were Archaea. The percentage occurrence of bacteria belonging to the phylum Proteobacteria, which are linked to the biotransformation of recalcitrant contaminants in wastewaters (24), predominates and correlated with respect to ecological variables like process operations (changing oxygen conditions), influent wastewater characteristics or geographical location (25). *Comamonadaceae* (40%) was the dominant family in cultures perturbed with 0.25 cycles/hr in high-DO regime while a higher abundance of *Methylophilaceae* (25%) was found in 1 and 2 cycles/hr treated cultures at high-DO range (Figure S2 in the supplementary information). Cultures exposed to 0.16 cycles/hr in low-DO regime showed a higher abundance of *Rhodocyclaceae* (15%). *Flavobacteriaceae* (10-12%) and *Pseudomonadaceae* (14%) were dominant in cultures perturbed with 0.25 and 0.5 cycles/hr at low-DO regime. The non-perturbed constant high and low-DO regime subjected cultures resulted in higher abundances (30-45%) for *Comamonadaceae*. dbRDA constrained ordination based on Bray-Curtis similarity was conducted to determine the extent to which bacterial community variation at family level can be explained by DO conditions (Figure 4). A linear relationship was confirmed between dependent variables (bacterial families, enzyme activity and residual OMP concentration) and changes in predictor variable (perturbed and non-perturbed DO conditions). The most important parameters contributing variation in bacterial community composition, assessed at the level of family was the presence of the bacterial families *Rhodocyclaceae, Syntrophaceae*, *Syntrophobacteraceae Sphingobacteriaceae*, *Xanthomonadaceae, Campylobacteriacea* and *Aeromonadaceae*, as well as laccase, lignin, horseradish peroxidase and cytochrome P450 activity, and OMPs (namely sulfamethoxazole, carbamazepine, naproxen and ibuprofen; Figure 4). PERMANOVA indicating the similarities and differences between the different DO treatments in terms of bacterial composition, enzyme activity and OMP removal efficiencies showed significant difference (*F* = 3.26; df = 10, 11; p = 0.001) between perturbed and non-perturbed cultures. Variation in bacterial community composition positively correlated with microbial enzyme activity (relate analysis, R = 0.46, p< 0.05) and OMP residuals (relate analysis, R = 0.54, p < 0.05).

**Figure 4.**
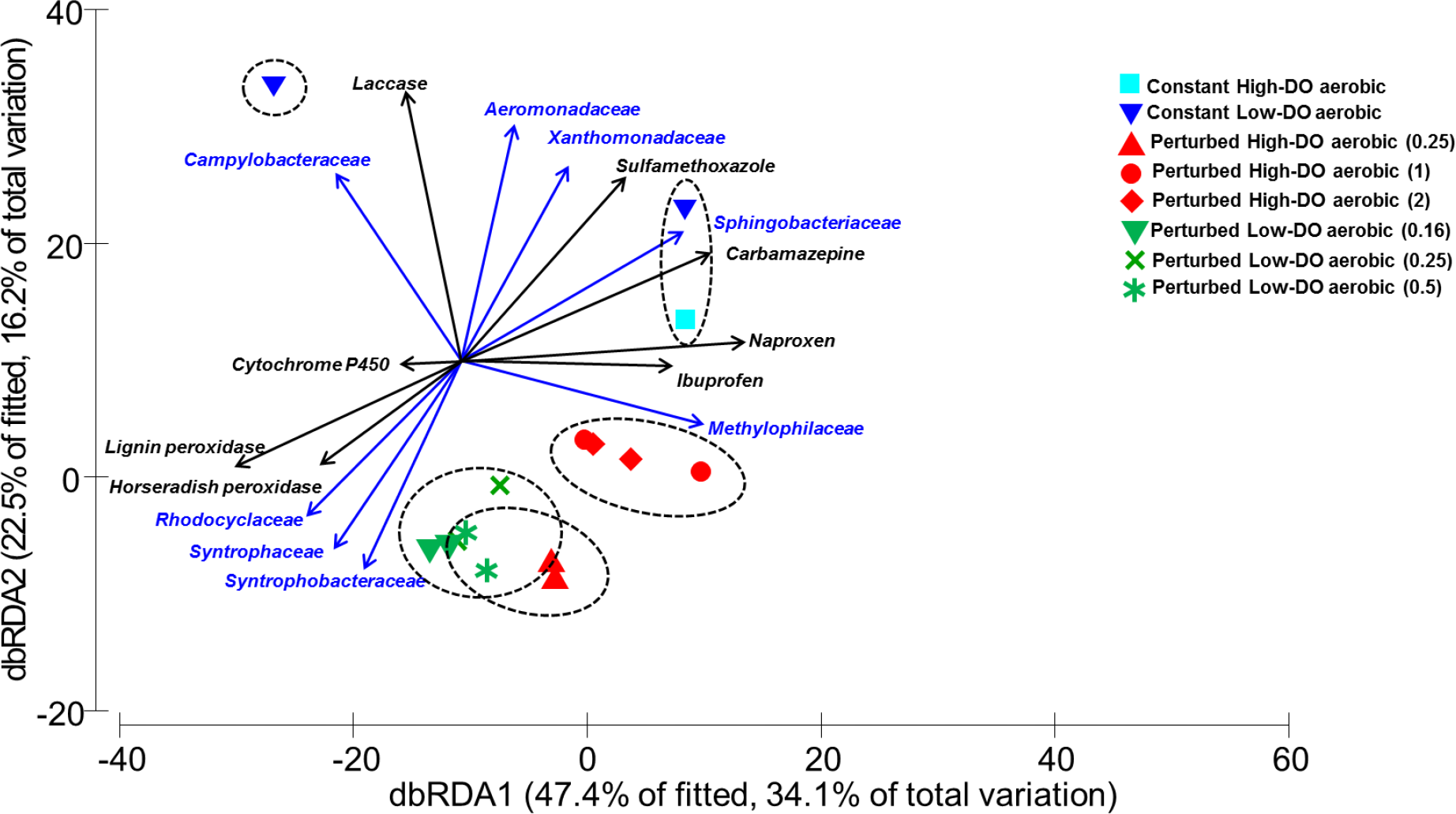
Variation in bacterial community composition (Bray-Curtis similarity) as revealed by distance-based redundancy analysis (dbRDA) of the data, illustrating relationships between bacterial families, enzyme activity and OMPs concentrations remaining in DO perturbed and non-perturbed control cultures at different frequencies. The dbRDA was constrained by the best fitting explanatory variables from multivariate multiple regression analysis (DISTLM) and vectors are shown for predictor variables explaining significant proportions of bacterial community variation (p = 0.001) at 97% OTU richness.

## Discussion

### Microbial Community Composition

Biological processes in waste treatment are manipulated by changing operation conditions (26) to promote specific eco-physiological characteristics of the microbial community to achieve the desired outcomes (25). Previously, the occurrence and abundance of Proteobacteria phyla has been linked to fluctuating concentrations of DO (27) and the effective biotransformation of pharmaceuticals (28). In the present study, perturbations of DO conditions have translated to relative differences in the microbial community composition. The core set of bacterial families included were *Commamonadaceae, Flavobacteriaceae, Rhodocyclaceae, Sphingobacteriaceae, Xanthomonadaceae, Burkholderiaceae and Pseudomonadaceae*, identified previously as major contributors to activated sludge treatment within wastewater treatment plants (WWTP) worldwide (29, 30). Genomic analysis has revealed that many common wastewater microorganisms have considerable potential to produce specialised enzymes and metabolites with OMP degradation potential (31–34). The core bacterial families observed in our study from the phylum Proteobacteria have been reported as organic micropollutant degraders in WWTPs treating pharmaceutical, petroleum refinery, domestic and agricultural wastewater (24). *Commamonadaceae* represented ~40% of the bacteria in both perturbed (0.25 cycles/hr, high-DO) and non-perturbed constant high-DO aeration regimes, followed by *Rhodocyclaceae* and *Syntropahceae* (15%) in 0.16 cycles/hr, low-DO perturbed conditions. These families can degrade aromatic compounds and denitrify industrial wastewaters (35). *Syntropahceae* dominated the anaerobic conditions within the 0.16 cycles/hr perturbed DO regime, consistent with other observations (36). However, *Flavobacteriaceae, Pseudomonadaceae* and *Sphingomonadaceae* (10-14%) families evident in 0.25 and 0.5 cycles/hr perturbed cultures at low-DO regime were identified as the core bacteria in activated sludge systems with floc-formation capabilities (37) and capable of co-metabolic biotransformation of ibuprofen (38), carbamazepine and sulfamethoxazole (39, 40). We found a significant correlation between microbial communities, enzyme activity, and OMP residual concentrations under different DO treated cultures. The bioremediation potential of laccases produced by *Pseudomonadaceae* and *Xanthomonadaceae* was successfully demonstrated (41). The cytochrome P450 system expressed in *Rhodococcus* spp. is known to degrade a range of wastewater-derived steroid hormones and toxic herbicides (42).

### Enzyme activity

Microbial enzymes are promising biocatalysts for OMP degradation (43); however, controlling the synthesis of poorly characterized enzymes to specifically enhance OMP treatment in complex wastewater communities has not been achieved. Oxygen is an obvious candidate for directed biocatalyst synthesis because it has a pronounced impact on the physiology and metabolism of microorganisms, including the induction of oxidoreductases demonstrated to degrade OMPs in pure enzyme assays. In this study, we found that cycling between aerobic and anaerobic phases stimulated the constant production of oxidoreductases (peroxidases, cytochromes and laccases) and native activated sludge enzymes such as hydrolases (beta-glucosidase and beta-glucosaminidase). The bacterial response to changing environmental oxygen was previously demonstrated by the activation of global regulatory systems – Arc BA, Fnr, OxyR and Sox RS (Figure 5) (44). The expression of peroxidases and cytochrome P450 in our study was approximately 1-2 orders of magnitude higher in the cultures perturbed with 0.25 and 0.5 cycles/hr at low-DO regime compared to 0.25 and 1 or 2 cycles/hr at high-DO regime treated cultures. Thus, exposing microbes to fluctuating aerobic-anaerobic phases induces the cellular enzymatic machinery to activate genetic adaptive strategies (e.g. *SoxRS* and *OxyR* mediated oxidative stress response systems). Gene activation leads to the synthesis of scavenging oxidoreductases, which provide a feedback control to lower the intracellular ROS concentrations (16, 45) (Figure 5). Increasing oxygen concentrations seem to result in the accumulation of significant amounts of intracellular ROS which overwhelm the cell with additional peroxidase expression. Increased levels of these enzymes are unable to completely counterbalance the toxic action of free oxygen radicals (46). However, with decreasing oxygen concentrations, anti-oxidative genes may remain induced and produce more active and efficient oxidoreductases (peroxidases and cytochromes) (47). The synthesis of beta-glucosidase is consistent with their involvement in organic matter degradation (48). Switching between aerobic and anaerobic conditions also yields bacterial metabolic reprogramming by regulating the activation of one component Fnr oxygen sensor and the two-component Arc system at the transcriptional level (44). The Fnr and Arc A systems exhibit maximal activity under microaerobic and anaerobic conditions (oxygen concentration ≤ 2.5% of total gaseous volume) leading to the mediation of metabolic responses, amplified production of redox signals, and more endogenous ROS generation (44, 49–51) (Figure 5). The regulation and expression of the oxidative genes by these regulatory systems allows a sensitive but robust transcriptional response over a wide range of oxygen concentrations. Proteomic analysis supported the correlation between maximum synthesis of cytochromes and peroxidases in the perturbed low-DO regime cultures. The synthesised oxido-reductases can simultaneously consume ROS as a cellular defence mechanism while oxidizing OMPs (52, 53). More studies would be needed to determine how ROS at cellular concentrations activates the expression of anti-oxidative genes and synthesis of oxidoreductases linked to the catalysis of OMPs.

**Figure 5.**
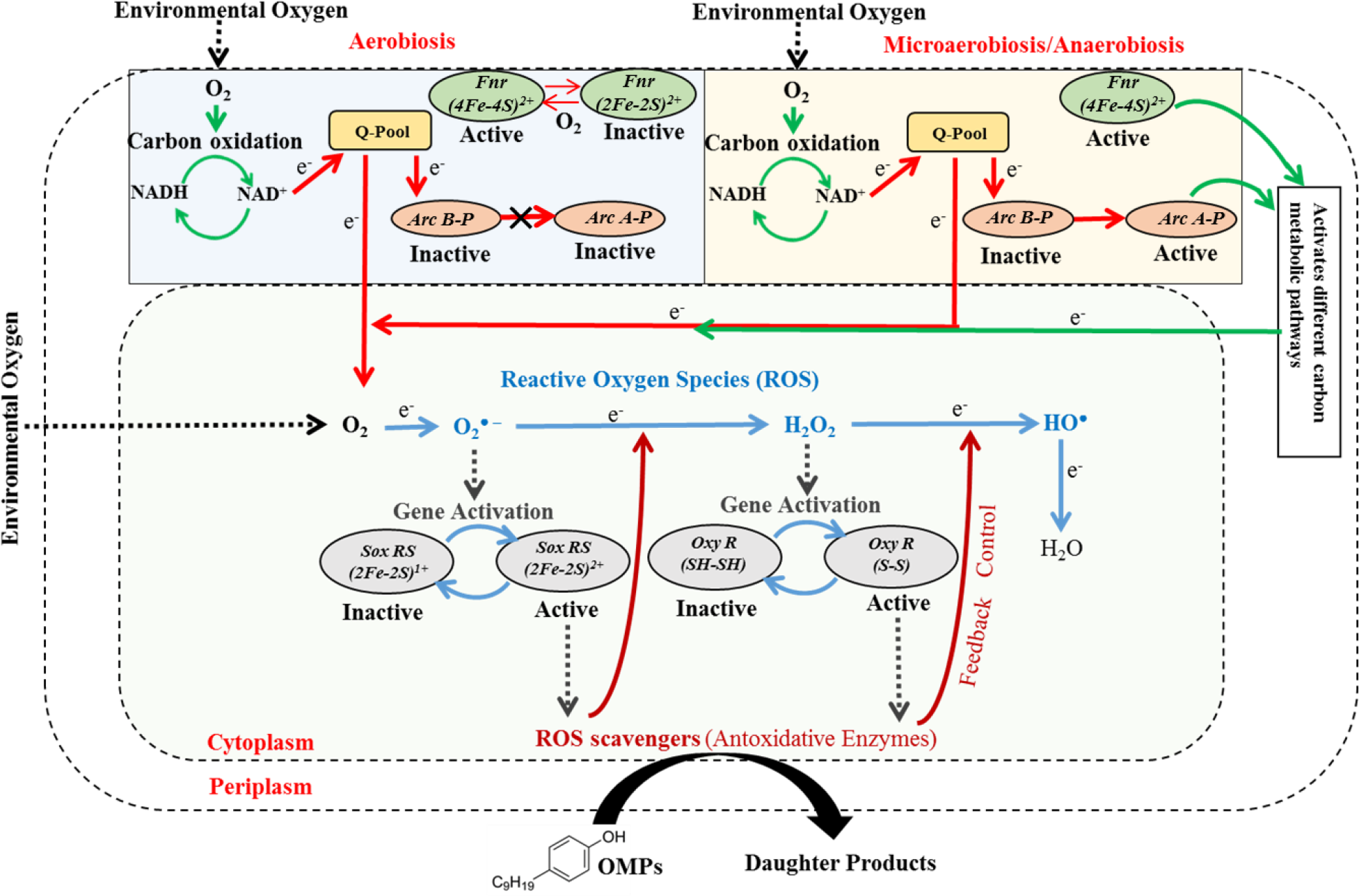
Regulation of enzyme production by the competition of ROS production and cellular respiration for O_2_. Under aerobiosis, NADH generated by carbon oxidation is oxidised to NAD^+^ and the electrons are passed on to oxygen by quinone pool (Q-pool). The autophosphorylation of membrane integrated Arc B is inhibited by oxidised quinone. Microaerobiosis/ Anaerobiosis causes a decrease in the level of oxidised quinones due to more production of NADH, allowing the transphosphorylation of Arc A-P. Arc A-P then coordinates the cellular response to microaerobiosis. Fnr is a direct sensor of oxygen availability. It is in a monomeric activated state with the incorporation of (4Fe-4S)^2+^ cluster. Under aerobic conditions, the active Fnr (4Fe-4S)^2+^ cluster gets converted to inactive (2Fe-2S)^2+^, thus, inhibiting the gene regulation while microaerobic/ anaerobic conditions block the Fnr at active (4Fe-4S)^2+^ form. Both Fnr and Arc AB regulates the cell metabolic pathways to generate redox signals which transfers electrons to the oxygen and generates the reactive oxygen species inside the cell. Another oxidative stress regulator (Sox RS and Oxy R) regulates the endogenous free radical concentrations by synthesis of antioxidative enzymes which simultaneously oxidise the organic micropollutants.

### Organic micropollutant removal

The efficiency of OMP biotransformation is linked to environmental conditions that enhance the expression of oxidoreductases by microbes (54). Such outcomes are consistent with results of this study where enhanced oxidoreductase production and OMP removal was observed in DO perturbed cultures compared to non-perturbed controls and autoclaved biomass. Similarly, sequential induction of aerobic and anaerobic conditions during biological wastewater treatment can enhance OMP biotransformation (55). Sludge cultures exposed to perturbations (0.16, 0.25, 0.5, 1 and 2 cycles/hr) in both low and high-DO regime showed more OMPs removal compared to controls with constant oxygen supply (Figure 3).

These data indicate that synthesis of highly active and efficient oxidoreductases (cytochrome P450, lignin and horseradish peroxidase) and hydrolases (beta-glucosidase) as a result of induced oxidative stress is a mechanism to exploit for OMP attenuation (56). Changing the oxygen conditions can influence the redox potential of the cell, hence, resulting in endogenous ROS production. As previously described, the oxygen free radicals can act as co-substrates for the synthesis of oxidoreductases (peroxidases, laccases and cytochromes) while fortuitously oxidising nearby organic substrates (53, 56). For example, the synthesised lignin peroxidases are high redox peroxidases which use activated molecular oxygen (such as superoxides and hydrogen peroxides) for the hydroxylation and cleavage of aromatic rings (57–59). We suggest that non-specific oxidoreductases such as peroxidases, cytochrome P450 and laccases in the DO perturbed cultures are metabolising OMPs in the presence of energy molecules (NADH or NAD^+^) produced during cellular metabolism operated by different oxygen regulatory systems (53).

The capability of synthesised oxidoreductases for OMP biotransformation across different functional groups including carboxylic compounds (NPX and IBP), hydroxyl compounds (TT) and other functional groups (SMX, CBZ and ATZ) is demonstrated by their transformation in the DO perturbed cultures (53, 60, 61). The active haem and copper centres of peroxidases and laccases and their reactivity with the evaluated OMPs (having relatively small sizes being comprised of 1-2 benzene rings) explains their significant biotransformation. The lower OMP removal efficiency of non-perturbed cultures is consistent with the generation of excess endogenous ROS, which partially inactivates peroxidases and allows for spontaneous conversion back into the resting enzyme (59). To improve the biodegradability of otherwise recalcitrant pharmaceuticals, personal care products and other micropollutants, the key step is to induce microorganisms to produce degradative enzymes. The efficient biodegradation of acetaminophen, acetyl sulfamethoxazole, atenolol and naproxen by oxidoreductases has been reported (39, 62), and 48-99% enzymatic transformation of carbamazepine, a highly recalcitrant molecule, by cytochrome P450 was demonstrated (63). The easy implementation, cost-effectiveness and potential to degrade a wide range of OMPs and other pollutants of concern suggests that this mechanism should be further investigated and exploited for its treatment potential.

This study suggests that exposing wastewater microbial species to oscillating oxygen concentrations in different aeration regimes can affect bacterial species selection, induce *de novo* synthesis of oxidative biocatalysts and subsequently enhance OMP removals. The use of such dissolved oxygen perturbations implies new and underutilized mechanisms for wastewater treatment plants to use existing infrastructure to improve their treatment efficiencies for problematic organic contaminants (14). With a potential to catalyse OMP biotransformations, the use of induced oxidoreductases in wastewater treatment has just started to be investigated at the laboratory scale (64). In future, the biodegradation of OMPs may be achieved by improving the performance of existing wastewater treatment systems under conditions that provide directed environmental stresses that result in over-production of beneficial OMP-degrading biocatalysts.

## Materials and methods

Additional details of the materials and methods can be found in the supplementary information.

### Reactor operational parameters and sampling

Fed-batch experiments were performed with mixed liquor sludge from an oxidation pond near the stables and milking areas of a dairy farm in Alfriston, New Zealand. Sludge was diluted to 3 g/L volatile suspended solids (VSS) with distilled water to a total volume of 1L. The bioreactors were continuously fed with ten times concentrated 100 ml synthetic wastewater (400 mg/L of COD as acetate and 60 mg-N/L ammonium as a nitrogen source), also containing 0.1 mg/L each of six model OMPs (detailed in the supplementary information) provided at a constant feed rate of 0.034 mL/min. for 48 hours. Oxidative stress was imposed on cultures by iteratively exposing them to ON-OFF cycles of an oxygen enriched air supply during the cultivation period. Additionally, constant (non-perturbed) high and low-DO conditions and autoclaved (dead) biomass were tested as controls. Each condition was tested in duplicate with sampling at the end of the cycle.

### Enzymes activity assays

The activity of oxidoreductases in culture biomass samples was determined spectrophotometrically by measuring the degradation (oxidation and hydrolysis) of various chromogenic substrates used as surrogate xenobiotics (Table S1 in the supplementary information). In brief, 20 ml aliquots of microbial culture samples were centrifuged at 18,078 x g for 15 min. at −4°C in 50 ml falcon tubes. The pellets from each tube were resuspended in either 10 ml acetate buffer or 10 ml phosphate buffer (detailed in supporting information). Each well of a 96 well microplate was filled with 100 μl aliquots of the respective buffers, 100 μl chromogenic dye and 100 μl resuspended biomass. Dyes and samples resuspended in double the amount of buffer served as controls. The prepared plates were mixed and incubated at 30°C for 1 hour. To start reactions of the sample enzymes with assay dyes, 10 μl of 0.3% H_2_O_2_ at 30% was added to the Methylene Blue, Azure B, L-DOPA and ABTS dye wells and 10 μl of 1M NaOH added to the para-nitrophenol dye wells (to stop the reaction) and vortex mixed. Changes in absorbance caused by chromogenic reactions were read on a Victor X3 Multimode Plate Reader (PerkinElmer, USA) at different wavelengths (refer to supporting information).

### Protein extraction and identification

Protein extraction was performed by cell lysis followed by precipitation of proteins and separation of low and high stringency protein fractions by 1D SDS PAGE. Briefly, 30 ml of sludge sample was pelleted and washed with 50 ml of 0.9% sodium chloride and spun down at 20,200 xg for 20 min at −4°C. The pellet was then washed again in 40 ml Tris-HCl (pH-7) and centrifuged at 20,200 xg for 20 min at −4°C to get the final pellet, which was resuspended in sample buffer, pulse-vortexed and, then placed on ice for 2 h with regular mixing at 15 min intervals. Samples were sonicated for 15 s (six rounds on ice) and centrifuged at 20,200 xg for 3 min at 4°C. The supernatant was mixed with trichloroacetic acid (100% w/v) and centrifuged at 23,400 xg for 30 min at −4°C after which the pellet was washed with 5 ml cold acetone twice. The final pellet was heat dried and re-suspended in 400 μl low stringency buffer (see supplementary information) to generate a low stringency fraction (LSF) in the supernatant, followed by centrifugation, then extracting this pellet in 400 μl high stringency buffer to generate a high stringency fraction (HSF). Proteins in both fractions were quantified spectrophotometrically by fluorescence. Crude separation of LSF and HSF proteins was achieved using 1D SDS PAGE. Individual bands were excised with a sharp razor blade and placed into low-binding, siliconized microcentrifuge tubes for destaining reduction, alkylation and trypsinolysis. Finally, 5 μL of the generated tryptic peptides of the microbial proteins were injected onto a SCIEX 6600 triple TOF mass spectrometer. Protein identification was carried out by comparing the obtained peptide sequences against those of the UniProt database. Additional methods and details can be found in the Supplementary Information.

### Extraction and analysis of organic micropollutants (OMPs)

OMPs at 0.1 mg each per L were prepared from a 1 g each per L stock solution in methanol and fed continuously at a feed rate of 0.034 mL/min to the bioreactor cultures for 48 h. Residual OMPs were extracted from the cultures using solid phase extraction (SPE) on OASIS HLB cartridges (Waters, Milford, MA, USA) following the method described by (65). Briefly, 200 mL of aqueous sludge culture was collected at the end of the cultivation cycle and centrifuged to remove suspended particles (20,4000 xg for 20 min at −4°C). The supernatant was passed through the cartridges (pre-conditioned with 5 mL each of tert-methyl butyl ether, methanol (Merck; laboratory grade purity) and deionised water) and washed with water. The OMPs were extracted in methanol (10 mL) under a vacuum system.

OMP analysis was performed by liquid chromatography coupled with mass spectrometry (LC-MS) using a Shimadzu 2020 Series LC-MS (Shimadzu, Japan) with an Agilent ZORBAX Eclipse Plus C18 column (2.1 mm × 100 mm, particle size 1.8 μm, Agilent Technologies, Germany) following EPA Method 1694 (66). A binary gradient system of mobile phase A, 0.1% formic acid in deionised water, and mobile phase B, 0.1% formic acid in acetonitrile were used to separate analytes in positive ESI mode, while 5 mM ammonium acetate, pH-5.5 (mobile phase A) and methanol (mobile phase B) were used for analysis in negative ESI mode. The solvent gradient programme for positive ESI mode was as follows: 10% B to 60% B in 24 min and then maintained at 100% B for 30 min. For negative ESI mode, the gradient started with 40% B, and was increased linearly to 100% B over 13.5 min, and then maintained at 40% B for 17 min. The flow rate was 0.2 ml/min. in both modes and the injection volume was set to 2 μl and 10 μl for +/− polarity modes, respectively. Limit of detection and limit of quantification were determined using signal/noise ratios of 3 and 10 respectively.

### Microbial DNA isolation and bacterial species identification

A PowerSoil DNA isolation kit (MoBio, Carlsbad, USA) was used for the isolation of bacterial total genomic DNA extracted from sludge samples (1 mL) following the manufacturer’s protocol. Bacterial community composition was characterised by amplifying and sequencing bacterial 16S ribosomal RNA (rRNA) gene fragments with the universal 16S PCR forward primer (5’-**TCGTCGGCAGCGTCAGATGTGTATAAGAGACAG**CCTACGGGNGGCWGCAG-3’) and 16S PCR reverse primer (5’ **GTCTCGTGGGCTCGGAGATGTGTATAAGAGACAG**GACTACHVGGGTATCTAA TCC-3’) following a standard protocol (67); nucleotide bases in bold are Illumina overhang adapter sequences for high throughput sequencing. Amplified PCR products were purified using an AMPure XP beads kit (Beckman Coulter Inc.) and concentrations recorded using a Qubit^®^ dsDNA HS Assay Kit (Life Technologies) according to the manufacturer’s instructions and then sequenced using an Illumina MiSeq machine. The resulting paired-end read DNA sequence data were merged and quality filtered using the USEARCH sequence analysis tool (68). Data were dereplicated so that only one copy of each sequence was reported. Sequence data were then checked for chimeric sequences and clustered into groups of operational taxonomic units based on a sequence identity threshold equal to or greater than 97% (thereafter referred to as 97% OTUs) using the clustering pipeline UPARSE (68) in QIIME v.1.6.0, as described in Bates et al. (69). Prokaryote phylotypes were classified to their corresponding taxonomy by implementing the RDP classifier routine (70) in QIIME v. 1.6.0 (71) to interrogate the Greengenes 13˙8 database (72). All sequences of chloroplast and mitochondrial DNA were removed. Finally, DNA sequence data were rarefied to a depth of 5,600 randomly selected reads per sample and two samples per treatment to achieve a standard number of sequencing reads across all samples.

### Statistical analyses

Enzyme activities were plotted using the heat map function within the R package ‘gplots’. Similarities in bacterial community composition, enzyme activity and OMP remaining in cultures were investigated using multivariate statistical analyses. Bray-Curtis distance matrices of relative bacterial abundance, enzyme synthesis and OMP residual concentrations were calculated and significant differences across all sample groups assessed using permutational multivariate analysis of variance (PERMANOVA). The RELATE function within the PRIMER package was used to calculate the Spearman rank correlation between data matrices constructed from the bacterial community, enzyme activity and OMP residual concentrations. All multivariate statistical analyses were performed in PRIMER 6 (Plymouth Marine Laboratory). Significant differences in OMP removal and residual concentrations were calculated by GraphPad Prism 7 using one-way ANOVA and post-hoc Tukey tests.

## Acknowledgements

The study was funded by FRDF grant 3707510/9572 to Singhal, Lear and Greenwood from the Faculty of Engineering at the University of Auckland. Bains is the recipient of a University of Auckland Doctoral Scholarship. Mabel Smith and Matthew Fung from the University of Alberta assisted with the development of analytical methods.

## References

1. Kim SD, Cho J, Kim IS, Vanderford BJ, Snyder SA (2007) Occurrence and removal of pharmaceuticals and endocrine disruptors in South Korean surface, drinking, and waste waters. Water Res 41:1013–1021.

2. Deblonde T, Cossu-Leguille C, Hartemann P (2011) Emerging pollutants in wastewater: A review of the literature. International J Hyg Environ Health 214:442–448.

3. Grandclément C, Seyssiecq I, Piram A, Wong-Wah-Chung P, Vanot G, Tiliacos N, Roche N, Doumenq P (2017) From the conventional biological wastewater treatment to hybrid processes, the evaluation of organic micropollutant removal: A review. Water Res 111:297–317.

4. Chang HS, Choo KH, Lee B, Choi SJ (2009) The methods of identification, analysis, and removal of endocrine disrupting compounds (EDCs) in water. J Hazard Mater 172:1–12.

5. Petrie B, Barden R, Kasprzyk-Hordern B (2014) A review on emerging contaminants in wastewaters and the environment: Current knowledge, understudied areas and recommendations for future monitoring. Water Res 72:3–27.

6. Luo Y, Guo W, Ngo HH, Nghiem LD, Hai FI, Zhang J, Liang S, Wang XC (2014) A review on the occurrence of micropollutants in the aquatic environment and their fate and removal during wastewater treatment. Sci Total Environ 473-474:619–641.

7. Graymore M, Stagnitti F, Allinson G (2001) Impacts of atrazine in aquatic ecosystems. Environ Int 26:483–495.

8. Cleuvers M (2003) Aquatic ecotoxicity of pharmaceuticals including the assessment of combination effects. Toxicol Lett 142:185–194.

9. Galus M, Jeyaranjaan J, Smith E, Li H, Metcalfe C, Wilson JY (2013) Chronic effects of exposure to a pharmaceutical mixture and municipal wastewater in zebrafish. Aquat Toxicol 132-133:212–222.

10. Joss A, Zabczynski S, Göbel A, Hoffmann B, Löffler D, McArdell CS, Ternes TA, Thomsen A, Siegrist H (2006) Biological degradation of pharmaceuticals in municipal wastewater treatment: Proposing a classification scheme. Water Res 40:1686–1696.

11. Stackelberg PE, Gibs J, Furlong ET, Meyer MT, Zaugg SD, Lippincott RL (2007) Efficiency of conventional drinking-water-treatment processes in removal of pharmaceuticals and other organic compounds. Sci Total Environ 377:255–272

12. Carballa M, Omil F, Lema JM, Llompart M, Garcı́a-Jares C, Rodrı́guez I, Gómez M, Ternes T (2004) Behavior of pharmaceuticals, cosmetics and hormones in a sewage treatment plant. Water Res 38:2918–2926.

13. Zorita S, Mårtensson L, Mathiasson L (2009) Occurrence and removal of pharmaceuticals in a municipal sewage treatment system in the south of Sweden. Sci Total Environ 407:2760–2770.

14. Falås P, Wick A, Castronovo S, Habermacher J, Ternes TA, Joss A (2016) Tracing the limits of organic micropollutant removal in biological wastewater treatment. Water Res 95:240–249.

15. McAdams HH, Srinivasan B, Arkin AP (2004) The evolution of genetic regulatory systems in bacteria. Nat Rev Genet 5:169–178.

16. Imlay JA (2008) Cellular Defenses against Superoxide and Hydrogen Peroxide. Annu Rev of Biochem 77:755–776.

17. Kaur A, Van PT, Busch CR, Robinson CK, Pan M, Pang WL, Reiss DJ, DiRuggiero J, Baliga NS (2010) Coordination of frontline defense mechanisms under severe oxidative stress. Mol Syst Biol 6:1–16.

18. Imlay JA (2013) The molecular mechanisms and physiological consequences of oxidative stress: lessons from a model bacterium. Nat Rev Microbiol 11:443–454.

19. Rao MA, Scelza R, Acevedo F, Diez MC, Gianfreda L (2014) Enzymes as useful tools for environmental purposes. Chemosphere 107:145–162.

20. Kumar S (2010) Engineering cytochrome P450 biocatalysts for biotechnology, medicine and bioremediation. Expert Opin Drug Metab Toxicol 6:115–131.

21. Riva S (2006) Laccases: blue enzymes for green chemistry. Trends Biotechnol 24:219–226.

22. Torres E, Bustos-Jaimes I, Le Borgne S (2003) Potential use of oxidative enzymes for the detoxification of organic pollutants. Appl Catal B: Environmental 46:1–15.

23. Gianfreda L, Xu F, Bollag J-M (1999) Laccases: A Useful Group of Oxidoreductive Enzymes. Bioremed J 3:1–26.

24. Gonzalez-martinez A, Sihvonen M, Muñoz-palazon B, Mikola A, Vahala R (2018) Microbial ecology of full-scale wastewater treatment systems in the Polar Arctic Circle : Archaea, Bacteria and Fungi. Sci Rep 2:1–11.

25. Isazadeh S, Jauffur S, Frigon D (2016) Bacterial community assembly in activated sludge: mapping beta diversity across environmental variables. MicrobiologyOpen 5:1050–1060.

26. Pholchan MK, Baptista J de C, Davenport RJ, Curtis TP (2010) Systematic study of the effect of operating variables on reactor performance and microbial diversity in laboratory-scale activated sludge reactors. Water Res 44:1341–1352.

27. Ouyang E, Liu Y, Ouyang J, Wang X (2017) Effects of different wastewater characteristics and treatment techniques on the bacterial community structure in three pharmaceutical wastewater treatment systems. Environ Technol 0:1–13.

28. Zhao C, Xie H, Xu J, Xu X, Zhang J, Hu Z, Liu C, Liang S, Wang Q, Wang J (2015) Bacterial community variation and microbial mechanism of triclosan (TCS) removal by constructed wetlands with different types of plants. Sci Total Environ 505:633–639.

29. Decai J, Ping W, Zhihui B, Xinxin W, Hong P, Rong Q, Zhisheng Y, Guoqiang Z (2011) Analysis of bacterial community in bulking sludge using culture-dependent and - independent approaches. J Environ Sci 23:1880–1887.

30. Cydzik-kwiatkowska A, Zielin M (2016) Bacterial communities in full-scale wastewater treatment systems. World J Microbiol Biotechnol 32:66.

31. Keller NP, Turner G, Bennett JW (2005) Fungal secondary metabolism - From biochemistry to genomics. Nat Rev Microbiol 3:937–947.

32. Miller LD, Mosher JJ, Venkateswaran A, Yang ZK, Palumbo AV, Phelps TJ, Podar M, Schadt CW, Keller M (2010) Establishment and metabolic analysis of a model microbial community for understanding trophic and electron accepting interactions of subsurface anaerobic environments. BMC Microbiol 10. doi:10.1186/1471-2180-10-149.

33. Mora-Pale M, Sanchez-Rodriguez SP, Linhardt RJ, Dordick JS, Koffas MAG (2014) Biochemical strategies for enhancing the in vivo production of natural products with pharmaceutical potential. Curr Opin Biotechnol 25:86–94

34. Rutledge PJ, Challis GL (2015) Discovery of microbial natural products by activation of silent biosynthetic gene clusters. Nat Rev Microbiol 13:509–523.

35. Alexander L, Claudia S, Sebastian L, Andreas SW, Kilian S, Christian B, Angelika L, Michael W (2005) 16S rRNA Gene-Based Oligonucleotide Microarray for Environmental Monitoring of the Betaproteobacterial Order “ Rhodocyclales.” Appl Environ Microbiol 71:1373–1386.

36. Meyer DD, de Andrade PA, Durrer A, Andreote FD, Corção G, Brandelli A (2016) Bacterial communities involved in sulfur transformations in wastewater treatment plants. Appl Microbiol Biotechnol 100:10125–10135.

37. Xu S, Yao J, Ainiwaer M, Hong Y, Zhang Y (2018) Analysis of Bacterial Community Structure of Activated Sludge from Wastewater Treatment Plants in Winter. Biomed Res Int doi: 10.1155/2018/8278970.

38. Barra Caracciolo A, Topp E, Grenni P (2015) Pharmaceuticals in the environment: Biodegradation and effects on natural microbial communities. A review. J Pharm Biomed Anal 106:25–36.

39. Fischer K, Majewsky M (2014) Cometabolic degradation of organic wastewater micropollutants by activated sludge and sludge-inherent microorganisms. Appl Microbiol Biotechnol 98:6583–6597.

40. Tran NH, Urase T, Ngo HH, Hu J, Ong SL (2013) Insight into metabolic and cometabolic activities of autotrophic and heterotrophic microorganisms in the biodegradation of emerging trace organic contaminants. Bioresour Technol 146:721–731

41. Majeau JA, Brar SK, Tyagi RD (2010) Laccases for removal of recalcitrant and emerging pollutants. Bioresour Technol 101:2331–2350.

42. Malandain C, Fayolle-Guichard F, Vogel TM (2010) Cytochromes P450-mediated degradation of fuel oxygenates by environmental isolates. FEMS Microbiol Ecol 72:289–296.

43. Nataliya MS, George SK, Anastasia AB, Alexey AD, Sergey LK, Kseniya MK, Nataliya VM, Anna VK (2016) Microbial community structure of activated sludge in treatment plants with different wastewater compositions. Front Microbiol 7:1–15.

44. Levanon SS, San KY, Bennett GN (2005) Effect of oxygen on the Escherichia coli ArcA and FNR regulation systems and metabolic responses. Biotechnol Bioeng 89:556–564.

45. Green J, Paget MS (2004) Bacterial redox sensors. Nat Rev Microbiol 2:954–966.

46. Baez A, Shiloach J (2014) Effect of elevated oxygen concentration on bacteria, yeasts, and cells propagated for production of biological compounds. Microb Cell Fact 13:1–7.

47. Podkopaeva DA, Grabovich MY, Dubinina GA (2003) Oxidative stress and antioxidant cell protection systems in the microaerophilic bacterium Spirillum winogradskii. Microbiology 72:534–541.

48. Krah D, Ghattas AK, Wick A, Bröder K, Ternes TA (2016) Micropollutant degradation via extracted native enzymes from activated sludge. Water Res 95:348–360.

49. Bekker M, Alexeeva S, Laan W, Sawers G, Teixeira de Mattos J, Hellingwerf K (2010) The ArcBA two-component system of Escherichia coli is regulated by the redox state of both the ubiquinone and the menaquinone pool. J Bacteriol 192:746–754.

50. Guest JR, Spiro S (1990) FNR and its role in oxygen-regulated gene expression in Escherichia coli. FEMS Microbiol Lett 75:399–428.

51. Georgellis D, Kwon O, Lin ECC (2001) Quinones as the redox signal for the Arc two-component system of bacteria. Science 292:2314–2316.

52. Schieber M, Chandel NS (2014) ROS function in redox signaling and oxidative stress. Curr Biol 24:R453–R462.

53. Gonzalez-Gil L, Carballa M, Lema JM (2017) Cometabolic Enzymatic Transformation of Organic Micropollutants under Methanogenic Conditions. Environ Sci Technol 51:2963–2971.

54. Alneyadi AH, Rauf MA, Ashraf SS (2018) Oxidoreductases for the remediation of organic pollutants in water–a critical review. Crit Rev Biotechnol 38:971–988.

55. Singhal N, Perez-Garcia O (2016) Degrading Organic Micropollutants: The Next Challenge in the Evolution of Biological Wastewater Treatment Processes. Front Environ Sci 4:1–5.

56. Khatoon N, Jamal A, Ali MI (2017) Polymeric pollutant biodegradation through microbial oxidoreductase: A better strategy to safe environment. Int J Biol Macromol 105:9–16.

57. Wen X, Jia Y, Li J (2009) Degradation of tetracycline and oxytetracycline by crude lignin peroxidase prepared from Phanerochaete chrysosporium - A white rot fungus. Chemosphere 75:1003–1007.

58. Bilal M, Rasheed T, Iqbal HMN, Yan Y (2018) Peroxidases-assisted removal of environmentally-related hazardous pollutants with reference to the reaction mechanisms of industrial dyes. Sci Total Environ 644:1–13.

59. Hofrichter M, Ullrich R, Pecyna MJ, Liers C, Lundell T (2010) New and classic families of secreted fungal heme peroxidases. Appl Microbiol Biotechnol 87:871–897.

60. Kassotaki E, Buttiglieri G, Ferrando-Climent L, Rodriguez-Roda I, Pijuan M (2016) Enhanced sulfamethoxazole degradation through ammonia oxidizing bacteria co-metabolism and fate of transformation products. Water Res 94:111–119.

61. Ingram-Smith C, Gorrell A, Lawrence SH, Iyer P, Smith K, Ferry JG (2005) Characterization of the acetate binding pocket in the Methanosarcina thermophila acetate kinase. J Bacteriol 187:2386–2394.

62. Kunkel U, Radke M, Kunkel UWE, Radke M (2008) Biodegradation of Acidic Pharmaceuticals in Bed Sediments : Insight from a Laboratory Experiment Biodegradation of Acidic Pharmaceuticals in Bed Sediments : Insight from a Laboratory Experiment. Environ Sci Technol 42:7273–7279.

63. Golan-Rozen N, Chefetz B, Ben-Ari J, Geva J, Hadar Y (2011) Transformation of the recalcitrant pharmaceutical compound carbamazepine by pleurotus ostreatus: Role of cytochrome P450 monooxygenase and manganese peroxidase. Environ Sci Technol 45:6800–6805.

64. Lah L, Podobnik B, Novak M, Korošec B, Berne S, Vogelsang M, Kraševec N, Zupanec N, Stojan J, Bohlmann J, Komel R (2011) The versatility of the fungal cytochrome P450 monooxygenase system is instrumental in xenobiotic detoxification. Mol Microbiol 81:1374–1389.

65. Vanderford BJ, Pearson RA, Rexing DJ, Snyder SA (2003) Analysis of Endocrine Disruptors, Pharmaceuticals, and Personal Care Products in Water Using Liquid Chromatography / Tandem Mass Spectrometry. Anal Chem 75:6265–6274.

66. Ferrer I, Thurman M (2008) EPA Method 1694: Agilent’s 6410A LC/MS/MS Solution for Pharmaceuticals and Personal Care Products in Water, Soil, Sediment, and Biosolids by HPLC/MS/MS: Application Note (Center for Environmental Mass Spectrometry University of Colorado Civil, Environmental, and Architectural Engineering, Boulder, CO).

67. Klindworth A, Pruesse E, Schweer T, Peplies J, Quast C, Horn M, Glöckner FO (2013) Evaluation of general 16S ribosomal RNA gene PCR primers for classical and next-generation sequencing-based diversity studies. Nucleic Acids Res 41:1–11.

68. Edgar RC (2013) UPARSE : highly accurate OTU sequences from microbial amplicon reads. Nat Methods 10:996–998. doi:10.1038/nmeth.2604.

69. Bates ST, et al. (2014) Biogeographic patterns in below-ground diversity in New York City’s Central Park are similar to those observed globally. Proc Biol Sci 281:20141988.

70. Wang Q, Garrity GM, Tiedje JM, Cole JR, Al WET (2007) Naıve Bayesian Classifier for Rapid Assignment of rRNA Sequences into the New Bacterial Taxonomy ᰔ †. Appl. Environ. Microbial. 73:5261–5267.

71. Caporaso JG, et al. (2010) correspondEnce QIIME allows analysis of high-throughput community sequencing data Intensity normalization improves color calling in SOLiD sequencing. Nat Methods 7:335–336.

72. Mcdonald D, et al. (2011) An improved Greengenes taxonomy with explicit ranks for ecological and evolutionary analyses of bacteria and archaea. ISME J 6:610–618.

73. Bassin JP, Pronk M, Kraan R, Kleerebezem R, Loosdrecht MCM Van (2011) Ammonium adsorption in aerobic granular sludge, activated sludge and anammox granules. Water Res 45:5257–5265.

