## Supplement for "Induction of microbial oxidative stress as a new strategy to enhance the enzymatic degradation of organic micropollutants in wastewater"

*[1]Department of Civil and Environmental Engineering, University of Auckland, New Zealand;[2]School of Biological Sciences, University of Auckland, New Zealand;[3]Faculty of Medical and Health Sciences, University of Auckland, New Zealand;[4]Division of Sciences and Mathematics, University of Washington-Tacoma; [5]Department of Civil & Environmental Engineering, University of Washington; [6]Centre for Urban Waters, Tacoma, WA, USA.*

**Corresponding Author: Naresh Singhal. Department of Civil and Environmental Engineering, University of Auckland, Private Bag 92019, Auckland 1142, New Zealand. Phone: +64 9 923 4512; Fax: +64 9 373 7462;*

### SUPPLEMENTARY METHODS

#### Reactor operational parameters and sampling

The mixed liquor sludge from a dairy farm was washed three times by mixing and decanting 0.5 L of supernatant with distilled water (Milli-Q system, Millipore, Darmstadt, Germany) in volumetric cylinders to achieve volatile suspended solids (VSS) of 3 g-VSS/L. The synthetic wastewater feed consisted of sodium acetate trihydrate (63 mM), magnesium sulphate heptahydrate (3.6 mM), potassium chloride (4.7 mM), ammonium chloride (35.4 mM), di-potassium hydrogen phosphate (4.2 mM), potassium dihydrogen phosphate (2.1 mM) and 10 mL/L of a trace element solution (73), all purchased from Sigma Aldrich; purity ≥ 98% (St. Louis, MO, USA). The model OMPs were as follows: the veterinary and human antibiotics sulfamethoxazole (SMX) and tylosin (TYL); the pharmaceuticals carbamazepine (CBZ), ibuprofen (IBP) and naproxen (NPX) and the agrochemical atrazine (ATZ); all of purity ≥ 98% and purchased from Sigma Aldrich (St. Louis, MO, USA). The stock (1 g/L) for the model OMPs was prepared in methanol (Merck, Darmstadt, Germany). Oxygen (99%) was purchased from BOC (Auckland, New Zealand).

#### Enzyme activity assays

Oxidoreductases such as lignin peroxidase, horseradish peroxidase, laccase (derived from cultures of *Trametes versicolor*), beta-glucosidase (from *Aspergillus niger*) and cytochrome P450 (from human 3A4 isozyme microsomes), as well as the respective enzyme substrates (Methylene Blue, Azure B, L-DOPA (3,4-Dihydroxy-L-phenylalanine), ABTS (2,2'-Azino-bis-(3-ethylbenzothiazoline-6-sulfonic acid)), Sudan Orange G, pNP-A (4-nitrophenyl N-acetyl-β-D-glucosaminide), pNP-G (4-nitrophenyl- β-D-glucopyranoside), pNP-12 (4-nitrophenyl-dodecanoate), Indole and 4-AAP (4-Aminoantipyrine)) were purchased from Sigma Aldrich (St. Louis, MO, USA). Sodium acetate trihydrate, glacial acetic acid (≥ 99% purity), di-potassium hydrogen phosphate, potassium di-hydrogen phosphate, dextrose and magnesium chloride hexahydrate of ≥ 99% purity were also obtained from Sigma Aldrich and used to prepare enzyme buffers at pH-5 and pH-7 respectively. The targeted oxidoreductases and their colorimetric probes are illustrated in Table S1. Two different buffers: 50 mM acetate buffer (50 mM sodium acetate trihydrate adjusted to pH-5 with glacial acetic acid) and 100 mM phosphate buffer (80 mM di-potassium hydrogen phosphate, 20 mM potassium dihydrogen phosphate, 10 mM dextrose, 6 mM magnesium acetate adjusted to pH-7.4) were used for the enzyme assays.

**Protein extraction and identification**

The high purity (≥ 99% purity) chemicals sodium chloride, tris-HCl, urea, thiourea, CHAPS, EDTA, dithiothreitol, Pefabloc SC, Pefabloc protector, trichloroacetic acid, triton-X, IPG, 2X Laemmlli buffers, Coomassie blue dye, iodoacetamide, ammonium bicarbonate, and trichloroacetic acid were purchased from Sigma Aldrich (St. Louis, MO, USA). Protein extraction was conducted for 2 days starting with a cell lysis phase followed by precipitation of proteins and separation of low and high stringency protein fractions by 1D SDS PAGE. On the first day, 30 ml of sludge sample was centrifuged at 20,000 xg for 20 min. The pellets were washed in 50 ml of 0.9% sodium chloride and spun down at 20,000 xg for 20 min at 4°C. Pellets were washed in 40 ml Tris-HCl (pH 7) and again pelleted down at 20,000 xg for 20 min at 4°C. The final pellets were resuspended in sample buffer following the recipe: 7M urea, 2M thiourea, 4% (w/v) CHAPS, 10 mM Tris-1 mM EDTA, 50 mM dithiothreitol, 25 mM Pefabloc SC and 2 mM Pefabloc protector and pulse-vortexed then placed on ice for 2 hours with regular mixing at 15 min intervals. The samples were sonicated for 15 sec. (6 rounds on ice) and centrifuged at 20,000 xg for 3 min at 4°C. Trichloroacetic acid (TCA: 100% (w/v)) was added to the supernatant so that the TCA concentration came to 10-20% and the samples incubated at -20 °C overnight. On the second day, the samples were centrifuged at 20,000 xg for 30 min at -4°C and the resulting pellet was washed with 5 ml cold acetone twice. The final pellet was heat dried to drive off acetone and then re-suspended by vortex mixing for 2 h in 400 µl low stringency buffer, comprising: 9M urea, 1% (v/v) Triton X-100, 1% (v/v) IPG buffer, 0.5% (w/v) dithiothreitol and a trace of bromophenol blue. The re-suspended samples were centrifuged at 20,000 xg for 45 min. resulting in the generation of a low stringency fraction (LSF) in the supernatant. The remaining protein pellet was re-suspended by vortex mixing for 2 hours in 400 µl high stringency buffer, comprising: 7M urea, 2 M thiourea, 4% (w/v) CHAPS, 1% (v/v) IPG buffer, 2% (w/v) dithiothreitol and a trace of bromophenol blue. The re-suspended samples were again centrifuged at 20,000 xg for 45 min resulting in the generation of a high stringency fraction (HSF) in the supernatant. Both fractions were quantified spectrophotometrically by fluorescence. Provisional separation of proteins was achieved using 1D PAGE. Briefly, sample preparation was carried out by dissolving pellets in 10 µl of MilliQ water and 10 µl of 2 X Laemmlli buffer and heating the mixture at 96°C for 2-3 minutes. The mixture was then cooled, centrifuged briefly and loaded onto a 4-12% SDS-PAGE gel. After running both LSF and HSF fractions on the gel and staining with Coomassie blue, zonal bands in the range of 250-350 kDa were each excised with a sharp razor blade and placed into low-binding, siliconised microcentrifuge tubes for destaining reduction, alkylation and finally trypsinolysis. Then 5 µL of the generated tryptic peptides of the microbial proteins was injected on a SCIEX 6600 triple TOF mass spectrometer. Protein identification was done by comparing the obtained peptide sequences against those of the UniProt database.

**Microbial DNA isolation and bacterial species identification**

A PowerSoil DNA isolation kit (MoBio, Carlsbad, USA) was used for the isolation of bacterial total genomic DNA extracted from sludge samples (1 mL) following the manufacturer's protocol. All the extractions were performed in duplicate. Bacterial community composition was characterised by amplifying and sequencing a fragment of the bacterial 16S ribosomal RNA (rRNA) gene following a standard protocol (Illumina 2013). The V3 and V4 region of 16S rRNA genes were amplified from individual DNA extracts with the universal 16S Amplicon PCR Forward Primer (5'-**TCGTCGGCAGCGTCAGATGTGTATAAGAGACAG**CCTACGGGNGGCWGCAG-3') and 16S Amplicon PCR Reverse Primer (5'-**GTCTCGTGGGCTCGGAGATGTGTATAAGAGACAG**GACTACHVGGGTATCTAATCC-3') (67). These primers have been validated to provide good bacterial phylum coverage as they are also modified to include Illumina adapter overhang sequences (in bold) required for downstream DNA sequencing. DNA amplification was conducted as follows: (i) 94˚C for 3 min; (ii) 30 cycles of 94˚C for 30 sec, 55˚C for 30 sec, 72˚C for 30 sec; (iii) 72˚C for 5 min. Following amplification, PCR products were purified using the AMPure XP beads kit (Beckman Coulter Inc., Brea, CA, USA) according to the manufacturer's instructions. The concentrations of purified amplicons were finally measured and recorded using a Qubit® dsDNA HS Assay Kit (Life technologies, Carlsbad, CA, USA) and submitted to New Zealand Genomics Ltd for sequencing by Illumina MiSeq machine. The resulting paired-end read DNA sequence data were merged and quality filtered using the USEARCH sequence analysis tool (68). Data were dereplicated so that only one copy of each sequence was reported, and 'singleton' sequences represented by only one DNA sequence in the database were removed. Sequence data were then checked for chimeric sequences and clustered into groups of operational taxonomic units based on a sequence identity threshold equal to or greater than 97% (thereafter referred to as 97% OTUs) using the clustering pipeline UPARSE (68) in QIIME v.1.6.0 as described in (69). After that, prokaryote phylotypes were classified to their corresponding taxonomy by implementing the RDP classifier routine (70) in QIIME v. 1.6.0 (71) to interrogate the Greengenes 13˙8 database (72). All sequences of chloroplast and mitochondrial DNA were removed. Finally, DNA sequence data were rarefied to a depth of

5,600 randomly selected reads per sample and two samples per treatment to achieve a standard sequencing reads across all samples.

### 1. Oxidoreductases and substrate dyes

**Table S1**. Target oxidoreductases and the dyes used to detect their activity in respective buffers.

| Target enzyme | Enzyme substrate (Dye) | Solution buffer |
|---|---|---|
| Lignin peroxidase | Methylene Blue | acetate buffer |
| (LiP) | Azure B | acetate buffer |
| Horseradish peroxidase | L-DOPA | acetate buffer |
| (HRP) | ABTS | acetate buffer |
| Laccase | Sudan Orange | acetate buffer |
| (Lac) | ABTS | acetate buffer |
| β-glucosaminidase (β-glcNAc) | pNP-A | acetate buffer |
| β-glucosidase (β-glu) | pNP-G | acetate buffer |
| Cytochrome P450 | pNP-12 | phosphate buffer |
| (Cyp450 or CYPcam) | Indole | phosphate buffer |
| | 4-AAP | phosphate buffer |

### 2. OMP concentrations in the bioreactors and statistical analysis

**Table S2.** Residual concentrations (mg/L) of OMPs under constant non-perturbed and perturbed DO frequencies (0.16, 0.25, 0.5, 1 and 2 cycles/hr) in different aeration regimes (high and low-DO) measured by LC-MS. Statistical differences among the samples are indicated with p<0.05 performed on duplicate sample set (n = 2) as mean ± standard deviation.

| Compounds | Constant High Aerobic | Constant Low Aerobic | Perturbed High Aerobic (0.25) | Perturbed High Aerobic (1) | Perturbed High Aerobic (2) | Perturbed Low Aerobic (0.16) | Perturbed Low Aerobic (0.25) | Perturbed Low Aerobic (0.5) |
|---|---|---|---|---|---|---|---|---|
| **Sulfamethoxazole** | 0.12 ± 0.02 | 0.09 ± 0.03 | 0.02 ± 0.00 | 0.05 ± 0.00 | 0.04 ± 0.01 | 0.04 ± 0.00 | 0.07 ± 0.07 | 0.01 ± 0.00 |
| **Carbamazepine** | 0.06 ± 0.02 | 0.06 ± 0.00 | 0.04 ± 0.00 | 0.05 ± 0.01 | 0.04 ± 0.01 | 0.03 ± 0.00 | 0.05 ± 0.06 | 0.04 ± 0.00 |
| **Tylosin** | 0.07 ± 0.01 | 0.07 ± 0.01 | 0.06 ± 0.01 | 0.06 ± 0.01 | 0.05 ± 0.01 | 0.05 ± 0.00 | 0.1 ± 0.1 | 0.05 ± 0.01 |
| **Atrazine** | 0.05 ± 0.00 | 0.04 ± 0.01 | 0.03 ± 0.00 | 0.02 ± 0.00 | 0.03 ± 0.01 | 0.01 ± 0.00 | 0.04 ± 0.05 | 0.02 ± 0.00 |
| **Naproxen** | 0.08 ± 0.00 | 0.05 ± 0.00 | 0.04 ± 0.00 | 0.05 ± 0.00 | 0.03 ± 0.01 | 0.03 ± 0.02 | 0.03 ± 0.01 | 0.02 ± 0.00 |
| **Ibuprofen** | 0.12 ± 0.03 | 0.08 ± 0.00 | 0.04 ± 0.00 | 0.05 ± 0.01 | 0.04 ± 0.00 | 0.03 ± 0.02 | 0.04 ± 0.01 | 0.02 ± 0.00 |

**Table S3.** Quality control parameters (Recovery percentage, Limit of detection (LOD) and Limit of quantification (LOQ)) of OMP extraction and analysis methods used. The LOD (mg/L) and LOQ (mg/L) are represented as mean ± standard deviation.

| Compounds | Recovery (%) | Limit of Detection (LOD) | Limit of Quantification (LOQ) |
|---|---|---|---|
| **Sulfamethoxazole** | 95 ± 2.2 | 0.060 | 0.181 |
| **Carbamazepine** | 100 ± 5.6 | 0.152 | 0.461 |
| **Tylosin** | 54 ± 2.5 | 0.095 | 0.288 |
| **Atrazine** | 100 ± 1.7 | 0.010 | 0.031 |
| **Naproxen** | 80 ± 2.4 | 0.117 | 0.354 |
| **Ibuprofen** | 100 ± 7.9 | 0.049 | 0.150 |

#### 3. Figure S1. Removal efficiency of OMPs form bioreactor culture at different aeration regimes

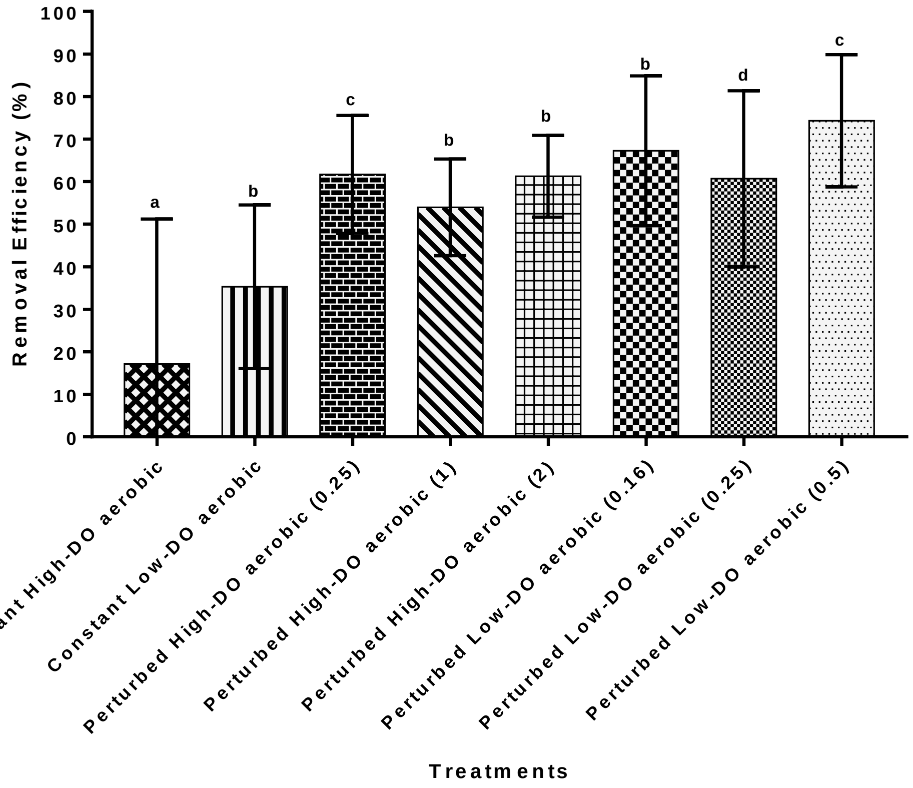


**Figure S1.** OMP removal efficiency was significantly increased with DO perturbation, as compared to within non-perturbed cultures ($p < 0.05$). Cultures under the perturbed low-DO aerobic regime (0.16, 0.25 and 0.5 cycles/hr frequencies) showed more OMP removal than cultures treated under perturbed high-DO aerobic (frequencies 1-2 cycles/hr) and non-perturbed constant high and low-DO aerobic regimes, respectively ($n = 2$). Different letters above bars denote significant differences between datasets according to post hoc Tukey tests at $p = 0.05$, treatments indicated with same letters are not statistically different.

### 4. Figure S2. Microbial Speciation with 16S rRNA gene sequencing

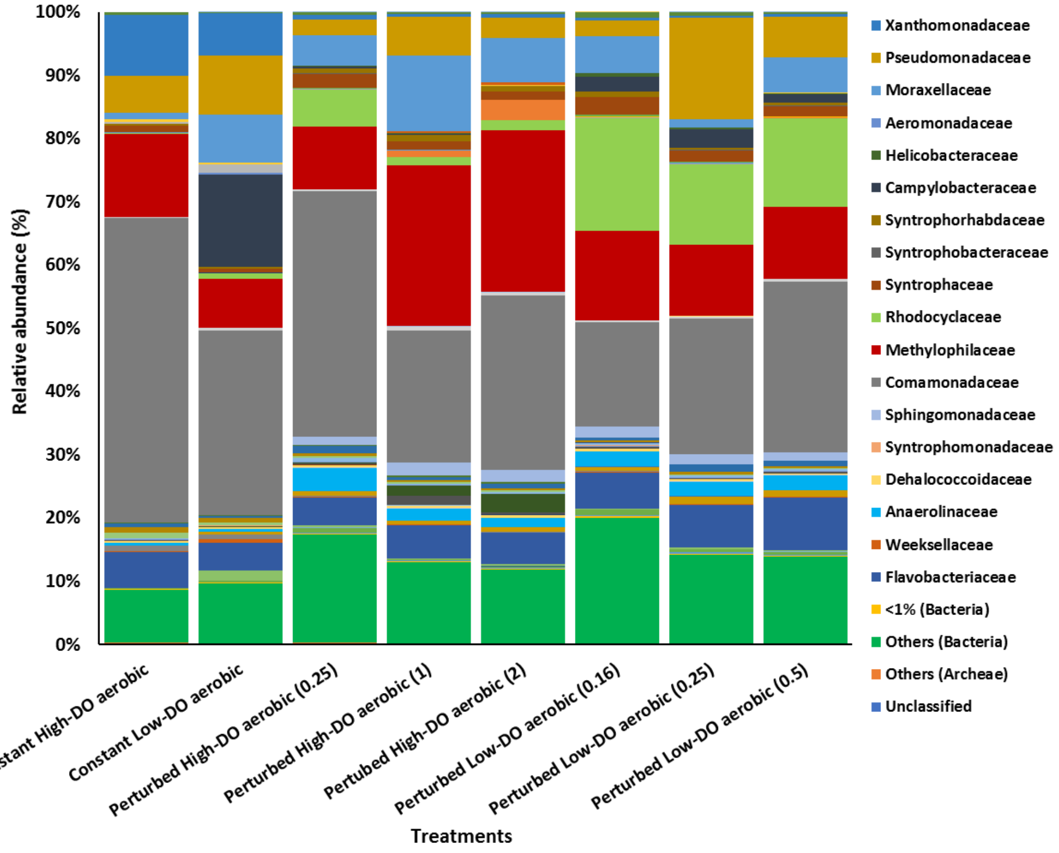


**Figure S2.** Composition of bacterial communities for taxa grouped at the family level in DO non-perturbed (constant) and perturbed) cultures. All analyses were done in duplicate (n = 2).
